# Post-saccadic face processing is modulated by pre-saccadic preview: Evidence from fixation-related potentials

**DOI:** 10.1101/610717

**Authors:** Antimo Buonocore, Olaf Dimigen, David Melcher

## Abstract

Humans actively sample their environment with saccadic eye movements to bring relevant information into high-acuity foveal vision. Despite being lower in resolution, peripheral information is also available prior to each saccade. How pre-saccadic extrafoveal preview of a visual object influences its post-saccadic processing is still an unanswered question. Here, we investigated this question by simultaneously recording behavior and fixation-related brain potentials while human subjects made saccades to face stimuli. We manipulated the relationship between pre-saccadic “previews” and post-saccadic images to explicitly isolate the influences of the former. Subjects performed a gender discrimination task on a newly foveated face under three preview conditions: phase-scrambled face, incongruent face (different identity from the foveated face), and congruent face (same identity). As expected, reaction times were faster after a congruent-face preview compared to the phase-scrambled and incongruent conditions. Importantly, a face preview (either incongruent or congruent) resulted in a strong reduction of post-saccadic neural responses. Specifically, we analyzed the classic face-selective N170 component at occipito-temporal EEG electrodes, which was still present in our experiments with active looking. We found that this component was strongly attenuated for face preview conditions compared to scrambled conditions. This large and long-lasting decrease in evoked activity is consistent with an active prediction mechanism influencing category-specific neural processing at the start of a new fixation. These findings constrain theories of visual stability and show that the extrafoveal preview methodology can be a useful tool to investigate its underlying mechanisms.

**Significance Statement:** Neural correlates of object recognition have traditionally been studied by flashing stimuli to the central visual field. This procedure differs in fundamental ways from natural vision, where viewers actively sample the environment with eye movements and also obtain a low-resolution preview of soon-to-be-fixated objects. Here we show that the N170, a classic electrophysiological marker of the structural processing of faces, also occurs during a more natural viewing condition but is massively reduced due to extrafoveal preprocessing (preview benefit). Our results therefore highlight the importance of peripheral vision during trans-saccadic processing in building a coherent and stable representation of the world around us.

## Introduction

In natural viewing, visual processing takes place primarily during periods of fixation, which are separated by fast and ballistic eye movements known as saccades. Unlike laboratory experiments, in which stimuli appear suddenly, the image that presents itself in the fovea at the beginning of each fixation is usually the result of a choice to fixate that item, typically based on a low-resolution peripheral preview of that object. Whether such a peripheral preview influences visual processing at the beginning of a new fixation, and how this might fit into different competing theories regarding why visual perception seems stable and continuous across saccades, remains an important question.

One view posits that the integration of pre- and post-saccadic information, at the level of neural mechanisms, involves a form of active prediction (Srinivasan et al., 1982; Rao and Ballard, 1999; Clark, 2013). The peripheral preview and the oculomotor plan for the next saccade might be combined to predict where the eye will land and what visual features will be present (for review, Melcher and Colby, 2008; Melcher, 2011). In the case of reading, a classic finding is the preview benefit effect in behavior (Rayner, 1975): when a word is visible in the periphery prior to the saccade, the subsequent fixation on the word is shorter compared to an invalid preview (Dimigen et al., 2012). When looking at fixation-related brain potentials (fERPs), Dimigen and colleagues have shown that such behavioral benefits are also associated with a reduction of the word-specific neural response, an effect termed the “preview positivity” (Dimigen et al., 2012; Kornrumpf et al., 2016).

Along these lines, several recent studies using fMRI have demonstrated a reduction in BOLD responses when a stimulus is consistent across the saccade (Dunkley et al., 2016; Zimmermann et al., 2016; Fairhall et al., 2017). For more complex images such as faces, there is behavioral evidence for an interaction across the saccade that can influence post-saccadic percepts (Melcher, 2005; Wolfe and Whitney, 2014). Recently, Edwards and colleagues (2017) showed that the decoding of post-saccadic EEG responses to faces was possible earlier when the preview of the target did not change during the execution of the saccade, suggesting the use of peripheral information.

An alternative view on active prediction focusses instead on the spatial shift of attention towards the peripheral stimulus. Prior to saccade execution, attention is directed towards the saccade target (Hoffman and Subramaniam, 1995; Deubel and Schneider, 1996; Zhao et al., 2012; Buonocore et al., 2017) and this attentional shift has been implicated in many theories of stable perception (Mathôt and Theeuwes, 2011; Melcher, 2011). A key idea here is that selective attention is immediately present at the beginning of the new fixation, leading to attentional facilitation of post-saccadic processing (for review, see Mathôt and Theeuwes, 2011). In contrast to active prediction, which typically results in a reduction in evoked responses, selective attention tends to amplify neural responses (for review, Thiele and Bellgrove, 2018). In the case of face stimuli, for example, selective attention enhances the evoked response to the stimulus (Mohamed et al., 2009; Sreenivasan et al., 2009; Churches et al., 2010). If overt attention shifts are similar to covert ones, then the post-saccadic fixation-related ERPs would be expected to be larger in amplitude when a preview is available, due to the target receiving attentional enhancement. Testing whether there is an increase in neural activity (due to attention) versus a reduction (due to prediction) has therefore been suggested to be a marker to differentiate between these two mechanisms (Kok et al., 2012; Spaak et al., 2016; de Lange et al., 2018).

The aim of the current study was to investigate whether a peripheral preview of a face image would influence the post-saccadic processing of that face and, if so, whether it would lead to an increase (attention) or reduction (prediction) in the neural response. We tested this hypothesis by measuring the effect of the pre-saccadic preview stimulus (either an intact or phase-scrambled face) on post-saccadic evoked response.

## Materials and Methods

### Participants

Seventeen volunteers (12 females) between 18 and 30 years of age (*M* = 23.6) participated in the study. All were free from neurological and visual impairments. The experiment was conducted in accordance with the Declaration of Helsinki (2008) and approved by the University of Trento Research Ethics Committee. All participants provided informed written consent and received a compensation of €10 per hour.

### Apparatus

Stimuli were presented on a 24-inch LED monitor (resolution: 1920×1080 pixels, subtending 39° × 24.7°) at a vertical refresh of 120 Hz. To reduce head movements, participants were seated with their head stabilized by a chin and forehead rest. The eyes were horizontally and vertically aligned with the center of the screen at a viewing distance of 63 cm. Eye movements were recorded with a video-based eye tracker (EyeLink 1000 with desktop mount; SR Research, Ontario, Canada) at a sampling rate of 1000 Hz (detection algorithm: pupil and corneal reflex; thresholds for saccade detection: 30 deg/s velocity and 9500 deg/s^2^ acceleration). A five point-calibration and validation of the eye tracker on a standard rectangular grid was run at the beginning of the experiment and whenever necessary during the experiment. Programs for stimulus presentation and data collection were written in MATLAB (MathWorks) using the Psychophysics Toolbox Version 3 (Brainard, 1997; Pelli, 1997) and Eyelink Toolbox extensions (Cornelissen et al., 2002). Participants’ manual responses were recorded on a standard keyboard.

The electroencephalogram (EEG) was recorded from 64 Ag/AgCl electrodes (Brain Products GmbH, Munich, Germany) placed at standard locations of the International 10-10 system. Signals were recorded with a time constant of 10 s and a high cutoff of 250 Hz, referenced online against the left mastoid, and digitized at a rate of 1000 Hz. The system was set up with a parallel port splitter so that trigger pulses were sent simultaneously to the EyeLink and EEG acquisition computers.

### Procedure

Participants were seated in a dimly lit room and then briefly familiarized with the task by the experimenter. Figure 1 illustrates the trial scheme. Participants started each trial by pressing the space bar while maintaining their gaze at a central fixation cross (0.5° wide, shown in white on a black background). One second after this button press, two circular placeholders (white rings, diameter 4°, line width 1 pixel) appeared to the left and right of the central fixation cross. Placeholders were centered at eccentricities of ±8° and indicated the positions of the upcoming preview stimuli. Once the eye tracker detected a stable fixation for 1000 ms within an area of 2° around the central fixation cross, the preview display was triggered. Depending on the condition, the preview display consisted either of two different scrambled faces (*scrambled-face preview* condition) or two different intact faces (*intact-face preview* condition) that appeared at the previous positions of the placeholders (see Fig. 1, panel “Preview”). After 500 ms of preview, the fixation cross changed its color and turned either green or red, thereby cueing the participant to execute a saccade towards the left or right stimulus, respectively (Fig. 1, panel “Saccade cue”). Participants were instructed to respond as quickly and accurately as possible to the cue with a single saccade.

**Figure 1.**
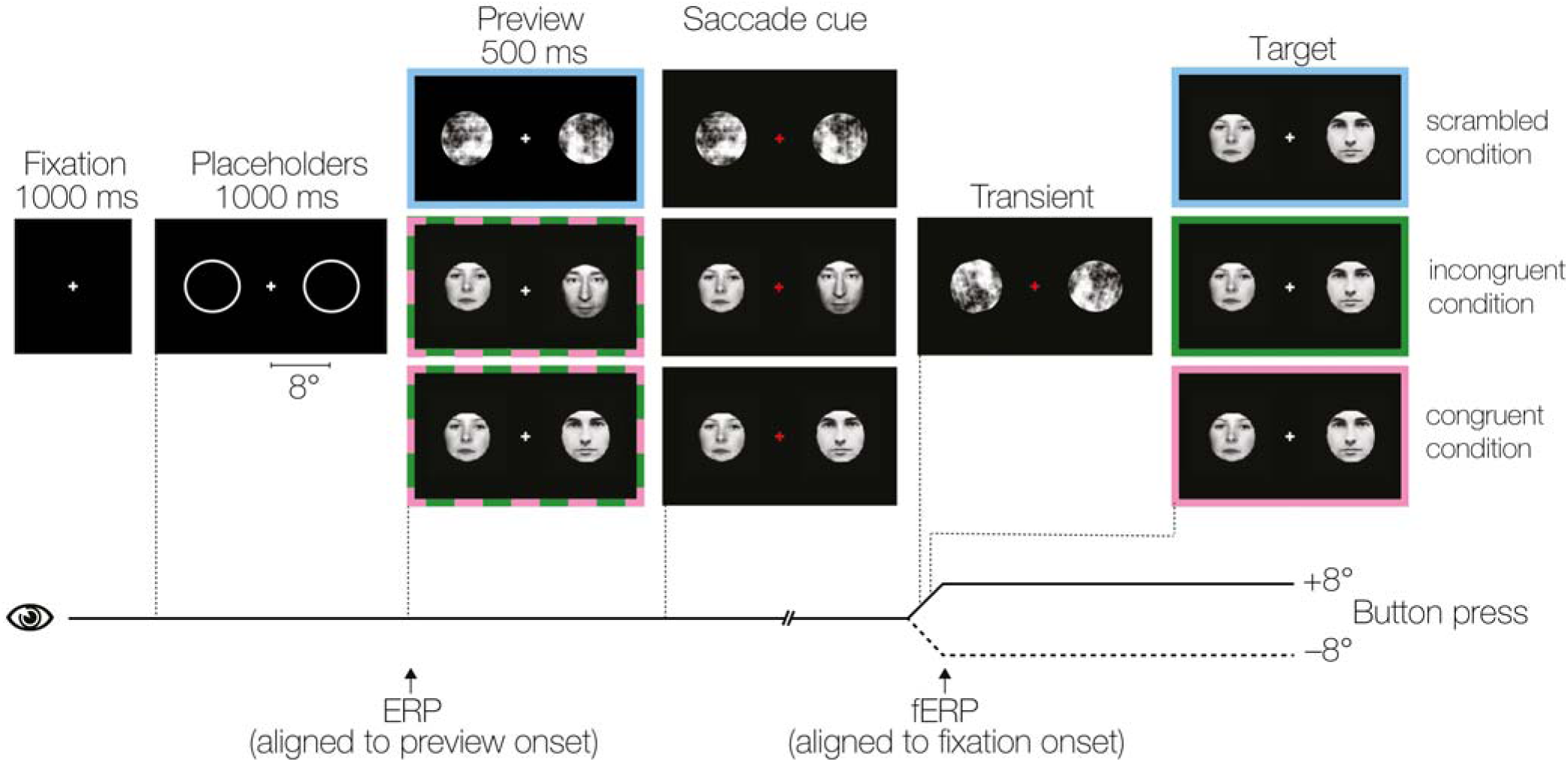
Trial scheme. At the beginning of each trial, participants fixated a central fixation cross 1000 ms. Afterwards, two placeholders appeared in the periphery at ±8 degree to the left and right of fixation (Placeholders panel). After 1000 ms, two preview stimuli appeared at the position of the placeholders for 500 ms (Preview panel). These stimuli could be either scrambled-faces (blue outline) or intact-faces or (dashed green-pink outline). After the preview interval, the central cross turned either green (left) or red (right), thereby cueing the participant to execute a saccade towards the left of right placeholder, respectively (Saccade cue panel). During the saccade, the preview was first changed into a scrambled image patch for one display cycle (8.3 ms) in order to introduce a peri-saccadic transient in all three conditions (Transient panel). Afterwards, the stimulus changed to the target face in all conditions (Target panel). The relationship between the preview stimulus and the target face yielded three conditions for the behavioral and fixation-related EEG analysis: a scrambled preview condition (blue outline), an incongruent preview condition (green outline; different face seen before and after saccade), and a congruent preview condition (pink outline; same face seen before and after saccade). Participants were asked to discriminate the gender (male/female) of the face visible after the saccade with a button press. Note that stimuli are not drawn to scale.

During the saccade, once gaze position crossed an invisible vertical boundary placed a distance of 1° from the fixation cross, a scrambled version of the preview face (that was always different from those shown as previews in the scrambled-face preview condition) was transiently presented for just a single display cycle (8.3 ms; see Fig. 1, panel “Transient”). The purpose of this gaze-contingent display change was to introduce an intra-saccadic visual transient in all experimental conditions, that is, also in the *congruent-face preview* condition in which the same face was presented before and after the saccade. After the transient was displayed, and still during the saccade, the preview stimulus always changed into an intact face (Fig. 1, panel “Target”). Participants then responded with a button press whether the face that they had landed on with their eyes was male or female. Responses were given with the index fingers of the left and right hand using two keyboard buttons.

The experimental design comprised three main conditions: *scrambled-face preview, incongruent-face preview*, and *congruent-face preview* (Fig. 1, panel “Preview”). Each condition comprised 160 trials, leading to a total of 480 trials. Conditions differed in terms of the stimulus shown before the saccade (preview stimulus). In the *scrambled-face preview* condition, the stimuli presented during the preview interval were scrambled faces. In contrast, in both the *incongruent- and congruent-face preview* conditions, the stimuli shown as previews were intact faces. After the saccade, participants always looked at a face as the target stimulus. This means that in the *scrambled-face preview* condition, the scrambled face shown as a preview changed into a face. In the *incongruent-face* preview condition, the target face shown after the saccade was different from the preview face seen before the saccade (in this condition, the face shown at the irrelevant screen location opposite the cued saccade direction remained the same). Finally, in the *congruent-face preview* condition, the target stimulus was identical to the face presented at this position before the saccade. The face seen after the saccade was equiprobably male and female and the gender of the target face was counterbalanced with the preview condition.

### Stimuli

Forty-two grayscale images were selected from the Nottingham face database (http://pics.stir.ac.uk/zips/nottingham.zip), each showing a frontal view of a face (21 female, 21 male) with a neutral facial expression. To standardize the images and to reduce differences between the genders, a black mask with a circular aperture was applied to each face to cover the external facial features (e.g. hair, see Fig. 1). The aperture was centered on the nose, spanned from the forehead to the chin, and subtended a diameter of 4° of visual angle at the viewing distance of 63 cm.

For each original face stimulus, we also generated a scrambled counterpart that was used as the pre-saccadic preview stimulus in the scrambled-face preview condition (see *Procedure*). For this purpose, we calculated the 2D Fourier transform of each face image and then added a matrix of random phase angles to the existing phase information of the image. We then performed an inverse Fourier transform, thereby preserving the original power spectrum of the image. The same circular aperture as for the intact faces was also applied to the scrambled images.

Finally, for each face image, we selected a second face stimulus that served as the saccade target in the condition with an incongruent preview as well as a third scrambled-face stimulus which was used as a transient during the saccade. Specifically, to control for low-level differences between the face stimuli shown before and after the saccade, we randomly selected for each image another face stimulus from the pool of 42 face images, such that their difference in average image luminance (estimated via their RGB grey values) was less than 4% (i.e. difference < 11 in 8-bit grey values) and not statistically significant (as confirmed by a one-way ANOVA). In addition, possible differences in image luminance between the stimulus shown before and after the saccade (see below) were also controlled by adding luminance as a predictor in the statistical analysis of the EEG (see section *Single-subject GLM*).

### Behavioral screening & analysis

In an initial analysis step, trials were screened for incorrect oculomotor behavior. Specifically, we removed all trials in which no saccade was executed towards either stimulus (0.1% of trials) or an eye blink occurred around the time of saccade execution (− 200 to 600 ms around saccade onset; 1.1%). Furthermore, we removed trials in which the eyes deviated from the central fixation cross by > 2° during the preview interval (1.9%), the saccadic reaction time (SRT) was extremely short (< 100 ms; 0.8%) or long (> 700 ms, 7.2%), saccade amplitude was extremely small (< 3°; 2.1%) or large (> 10°; 2.9%), or in which the saccade went in the wrong direction (6.0%). Finally, we excluded trials in which the saccade-contingent display change was triggered prematurely by drift movements or microsaccades during the preview interval (11.5%) or in which the main saccade to the target was followed by a secondary saccade larger than 3° within 150 ms or less (0.2%). After applying these behavioral criteria, two participants had to be excluded from the sample due to excessive trials loss (>60%), reducing the final sample to 15 participants.

Manual RTs and response accuracy in the gender discrimination task were then submitted to repeated-measures ANOVAs on the three-level factor *Preview*. For the analysis of the button presses, trials with an extreme manual RT (< 200 or > 1000 ms) were ignored as outliers. Furthermore, one participant was dropped from the manual RT analysis due to very slow manual RTs and therefore too few remaining trials.

### Electrophysiological data analysis

For the electrophysiological analysis, the EEG was first synchronized with the eye-tracking channels based on the shared trigger pulses using the EYE-EEG toolbox (Dimigen et al., 2011). The synchronized EEG was then downsampled to 500 Hz, bandpass-filtered from 0.1 to 40 Hz using EEGLAB’s (Delorme and Makeig, 2004) finite response filter (*pop_eegfiltnew.m*) with default settings, and digitally re-referenced to an average reference. In the next step, ocular EEG artifacts were removed using an optimized eye-tracker-guided variant of Infomax ICA in EEGLAB. To optimize the ICA decomposition and the suppression of the myogenic spike potential peaking at saccade onset (Keren et al., 2011), the ICA was trained on a copy of the data high pass-filtered at 2 Hz (Winkler et al., 2015) in which EEG sampling points occurring around saccade onsets (−20 to +10 ms) were overweighted (see Dimigen, 2018). The resulting unmixing weights computed on this high-pass filtered and optimized training data were then applied to the original unfiltered recording, and ocular components were automatically flagged using the eye tracker-guided procedure by Plöchl et al. (2012) with the saccade/fixation variance ratio threshold set to 1.1 (Plöchl et al., 2012; Dimigen, 2018).

Based on the trials with a correct oculomotor behavior, we then extracted two sets of 1000 ms long epochs (−300 to 700 ms) from the artifact-corrected EEG. The first set was cut around the onset of the preview stimuli on the screen (traditional ERP average). The second set was cut around the onset of the first fixation on the target face following the saccade (fERP average). To exclude segments with residual non-ocular artifacts, we removed all epochs containing peak-to-peak voltage differences > 120 µV in any channel (2.3% of ERP and 2.8% of fERP epochs). Epochs were then baseline-corrected by subtracting the mean channel voltages in the 200 ms interval before stimulus/fixation onset, respectively.

### Single-subject GLM (first-level analysis)

Stimulus- and fixation-related potentials were analyzed using a massive univariate model (Smith and Kutas, 2015a) in which a GLM was fitted on each electrode and time point separately using the *unfold* toolbox (Ehinger and Dimigen, 2018). Analysis of EEG data with massive univariate models has advantages in terms of higher sensitivity and unbiased data-driven analysis (Rousselet et al., 2011; Smith and Kutas, 2015a) and allows to control the effects of continuous covariates on the waveform. For ERPs, the model only contained the intercept term and one categorical predictor coding whether the preview stimuli consisted of two scrambled (0) or two intact faces (1). For the fERP analysis, the predictors in the regression model were a three-level categorial predictor coding the type of preview shown before the saccade (scrambled, incongruent, congruent) as well as two continuous linear covariates: saccade amplitude and the preview-target luminance difference. Saccade amplitude (in degrees of visual angle) was added in the model because the size of the incoming saccade has a well-established and strong influence on the amplitude of the post-saccadic neural response (e.g. Thickbroom et al., 1991; Dandekar et al., 2011). Including saccade amplitude as a nuisance variable in the model therefore controlled for the slight difference in incoming saccade amplitude (about 0.3°, see *Results*) between the preview conditions. In addition, we also found that the fERP was modulated by the difference in mean luminance between the stimulus shown as preview and the post-saccadic target. The mean luminance difference between both stimuli was therefore also included as a continuous covariate.

As a control analysis, we repeated our analysis of the fERP using a GLM-based linear deconvolution technique (also called continuous-time regression, Dandekar et al., 2011; Smith and Kutas, 2015b; Ehinger and Dimigen, 2018) that is also implemented in the *unfold* toolbox. In the current experiment, SRTs were about 30 ms longer for the scrambled-face preview than for the intact-face preview conditions (see *Results*). This means that the temporal overlap between the ERP evoked by the onset of the saccade cue (red/green fixation cross) and the fERP evoked at saccade offset differed systematically between conditions, potentially biasing the results. GLM-based deconvolution allows us to control this overlapping activity by modeling the response to both types of events (cue and fixation onset) in the same statistical model. However, since the results were virtually identical to those obtained with the simpler univariate model, we only report the results of the latter here.

### Group statistics (second-level analysis)

Second-level statistical analyses were performed using the threshold-free cluster enhancement method (TFCE, Smith and Nichols, 2009; Mensen and Khatami, 2013), a variant of a cluster-based permutation tests (Maris and Oostenveld, 2007) which controls for multiple testing across electrodes and time points without the need to define an arbitrary cluster-forming threshold. Analyses were run using the Matlab implementation of TFCE (http://github.com/Mensen/ept_TFCE-matlab) based on 2000 random permutations. For ERPs, we compared the response following an intact-face vs. scrambled-face preview. For fERPs, we used the ANOVA variant of the TFCE algorithm, followed up by Bonferroni-corrected pairwise comparisons between the three preview conditions, again using the TFCE method. For visualization of the TFCE results in Figures 2 and 4, *p*-values were thresholded at *p*<0.05, *p*<0.01, and *p*<0.005.

**Figure 2.**
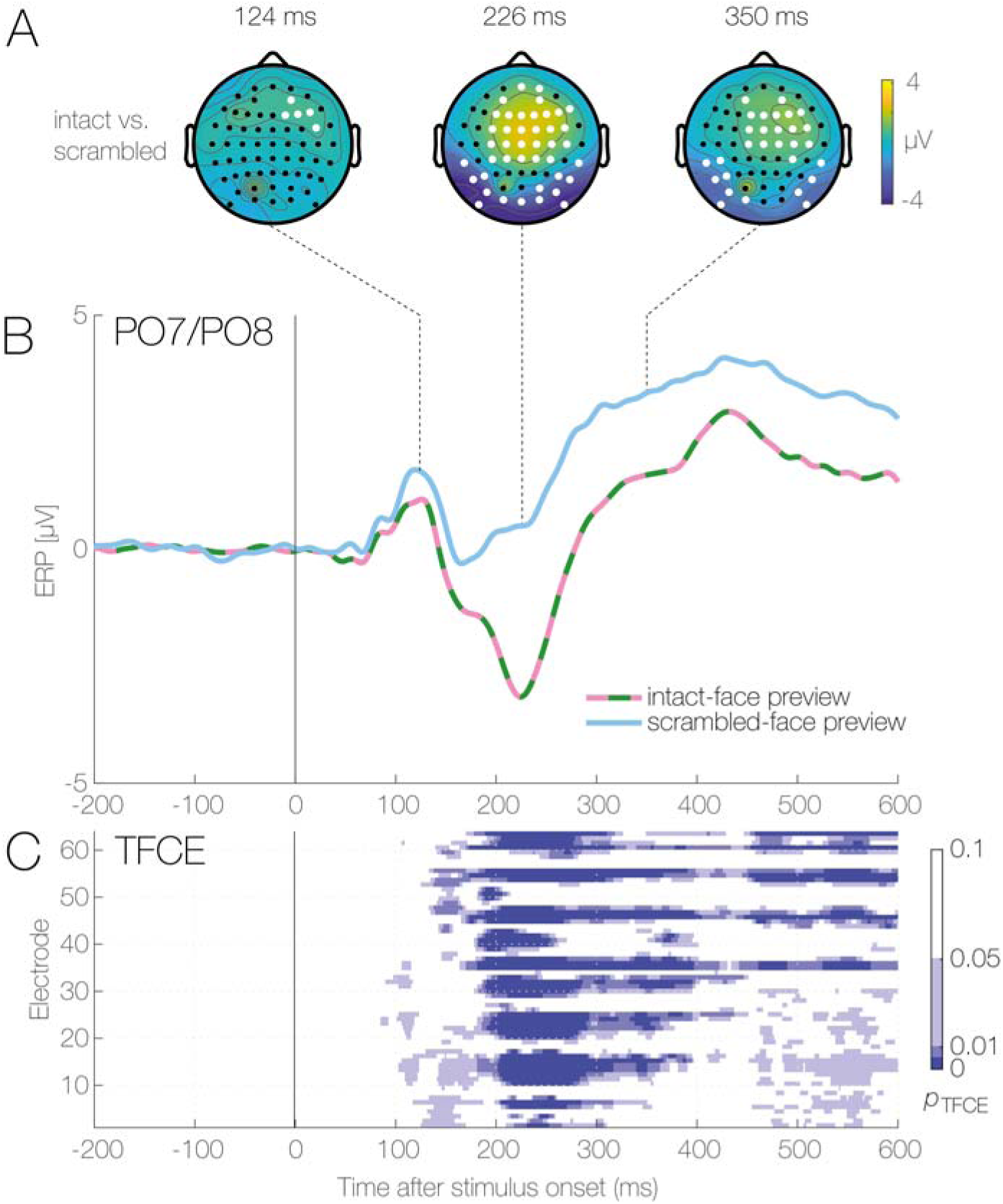
Event-related potentials aligned to the onset of the preview display. (A) Topographic difference maps between intact-face previews minus scrambled-face previews for three latencies after stimulus onset that represent the peak latencies of the P1, N1, and P300, respectively. White dots represent electrodes that show significant differences between the two preview conditions in the TFCE statistic at this latency. (B) Grand-mean stimulus-locked ERP, averaged over occipito-temporal electrodes PO7 and PO8 for intact-face previews (green-pink) and scrambled-face previews (blue). (C) Results of the TFCE statistic comparing face- and scrambled-face previews at all time points and channels. For visualization, *p*-values are thresholded at 0.05, 0.01, and 0.005 with different shades of blue.

## Results

In the following, we first report the neural response evoked by the onset of the preview stimuli (ERP to intact vs. scrambled faces). This is followed by an analysis of the behavior and fERP to the post-saccadic face stimulus.

### Preview stimulus onset: evoked response (ERP)

The goal of this analysis was to ensure that our stimuli were effective in eliciting typical face-related ERP components. Figure 2A shows the scalp-topographic difference maps of the difference between extrafoveal intact-face previews (i.e. two faces presented bilaterally at ±8° eccentricity) minus scrambled-face previews (two scrambled faces presented at ±8). Topographies are shown at three latencies after preview stimulus onset, corresponding to the peaks of the P1 (124 ms), N1 (226 ms) and P3 (350 ms) components. White dots in the scalp maps indicate electrodes which showed significant differences between intact- and scrambled-face previews at the given latency (in a pairwise TFCE-based *t*-test). The grand-mean waveforms in Figure 2B show the stimulus-ERP elicited by the onset of the bilateral preview display, averaged across two occipito-temporal electrodes over the left (PO7) and right hemisphere (PO8).

At the earlier latencies, during the P1 component, there was not yet a clear difference between the ERP responses for the two types of stimuli (intact-vs. scrambled-faces) beside a small cluster of activation at right frontocentral sites. However, in the following N1 time window, a strong bilateral negativity emerged at occipital-temporal electrode sites that was slightly larger over the right hemisphere, as typical for N170 face effects (Eimer, 2012). Over frontocentral sites, the posterior N170 effect was accompanied by a corresponding “vertex positive potential” (VPP); a broad positive potential generally taken to reflect the positive poles of the bilateral dipole pair giving rise to the occipito-temporal N170 (Eimer, 2012). These results clearly therefore show how the bilateral presentation of the face preview (dashed green-pink line) led to a markedly different evoked response than that of the scrambled-face images (blue line) (Fig. 2B); with faces eliciting a much more pronounced N170 component over occipital-temporal sites (Halgren et al., 2000; Hoshiyama et al., 2003; Deffke et al., 2007; Gao and Wilson, 2013). In contrast, only a weaker effect (i.e. smaller frontocentral cluster) of preview type was observed during the earlier P1 component (Fig. 2A). With a peak at around 226 ms, the N170 reached its peak about 50 ms later than typically observed (Bentin et al., 1996). A likely reason for this delayed N170 peak is that the two face stimuli were presented bilaterally in the extrafoveal visual field, rather than in the fovea. By looking at the full matrix of TFCE-corrected *p*-values depicted in Fig. 2C, it is clear how clusters of significant activation arose at around 160 ms after stimulus onset, both over antero-frontal areas and occipito-temporal electrodes. Although the difference between the intact-face and scrambled-face preview condition reached its maximum after 224 ms, this effect remained topographically stable and statistically significant throughout the entire stimulus-locked analysis period (i.e. until 600 ms after stimuli onset).

### Preview effects: behavioral results

Figure 3 summarizes behavioral performance in the task. A first finding is that saccadic reaction times (SRTs) were affected by the preview condition: SRTs were about 30 ms faster in trials with an intact compared to a scrambled-face preview (intact vs. scrambled: *t*(14) = −4.673; *p* < 0.0004) (Fig. 3A). The same pattern was also reflected in the saccade amplitudes, which were slightly larger (∼0.3°) when the preview was an intact rather than a scrambled face (Fig. 3B, *t*(14) = 8.259; *p* < 0.000001). This pattern of results indicates that seeing a possible target stimulus – that is, a face – in the periphery enhanced the preparation of the oculomotor response toward the target.

**Figure 3.**
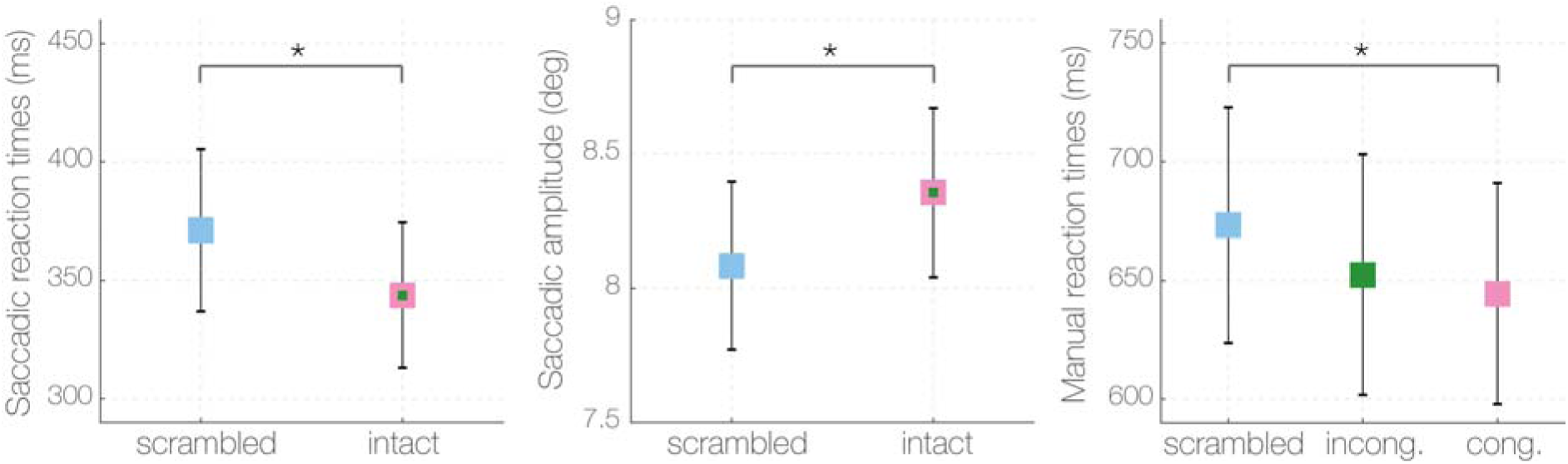
Behavioral results. Panels show the average (A) saccadic reaction time, (B) saccadic amplitude, and (C) manual RT for the scrambled-, incongruent-, and congruent-face preview condition, respectively. Asterisks denote *p* < 0.05. Error bars denote ±1 SEM.

For the gender discrimination task following the saccade, response accuracy was generally high (89% correct) and did not differ between preview conditions (*F*(2,26) = 0.475 *p* < 0.627). However, like SRTs, manual RTs for the button press depended strongly on the preview condition (main effect; *F*(2,26) = 8.535 *p* < 0.001) with numerically shorter RTs observed in the two conditions in which a congruent- or an incongruent face was shown as a preview compared to the scrambled-face condition (Figure 3, right panel). Bonferroni-corrected post-hoc tests confirmed that congruent face previews produced significantly shorter RTs than scrambled previews, *t*(13) = −3.802; *p* < 0.007. Importantly, this effect replicates the classic trans-saccadic preview benefit also observed with other types of stimuli, in particular words (Rayner, 1975; Dimigen et al., 2012). When the preview was an incongruent face, there was a only a statistical trend for faster RTs as compared to the scrambled-preview condition, *t*(13) = −2.546; *p* < 0.07. Manual RTs did not differ significantly between the congruent and incongruent preview condition.

Taken together, these results replicate a robust trans-saccadic benefit for previewed human faces compared to a non-informative scrambled-preview condition. Both the initial oculomotor response towards the peripheral face as well as the subsequent foveal processing of the facial features (necessary for the gender discrimination task) were significantly enhanced if the extrafoveal preview provided before the saccade was also a human face, supporting the hypothesis of preview facilitation for the processing of face stimuli.

### Preview effects: evoked response (fERP)

Main goal of the current study was to compare the fixation-related brain response elicited by the first direct fixation on the target face as a function of the extrafoveal information available during the *preceding* fixation: a scrambled face, a different person’s face, or the same face.

Figure 4 summarizes the fERP elicited by the first direct fixation on the target face, that is, after the end of the critical saccade. Top panels (Figure 4A) show the topographic difference maps for the three contrasts at the peaks of the P1 (106 ms), N1 (180 ms) and P3 (350 ms) component. The middle panel shows the corresponding fERP waveforms, averaged again across occipito-temporal electrodes PO7 and PO8. The bottom panel (Figure 4C) presents the corresponding statistical comparison (TFCE) between the congruent- and the scrambled-face preview conditions and between the incongruent- and the scrambled-face preview conditions (Figure 4D).

**Figure 4.**
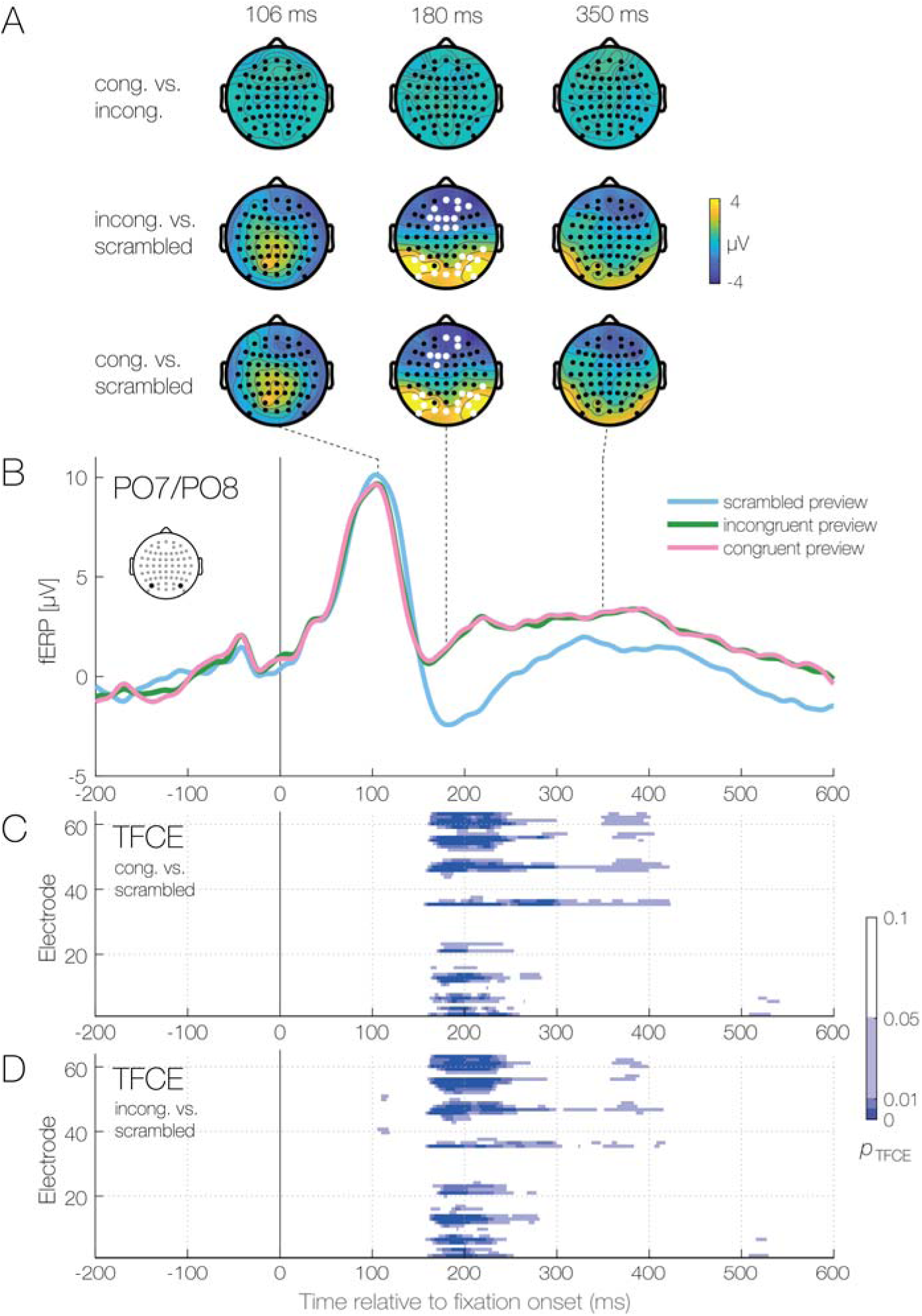
Fixation-related potentials (fERP). (A) Topographic difference maps for the difference between the congruent vs. incongruent (top row), incongruent vs. scrambled (middle row) and congruent vs. scrambled preview condition (bottom row) at three latencies after fixation onset on the target face. The latencies correspond to the P1, N1, and P300, respectively. (B) Grand-mean fERP averaged across occipito-temporal electrodes PO7 and PO8 for the scrambled (blue), incongruent (green) and congruent face preview condition (blue). Note that the three conditions only differ in terms of the stimulus seen before the saccade, the target face fixated at time zero of this plot was the same in all three conditions. (C-D) TFCE results for the pairwise comparison between the congruent- and scrambled-face preview condition and the incongruent- and scrambled-face preview condition respectively.

The first interesting observation is that when contrasting the activity following a congruent- as compared to an incongruent-face preview (Fig. 4A, top row), there was no sign of a significant difference across the entire scalp for any of the main components. In the second and third rows of Fig. 4A, we contrasted the activity of the incongruent- or congruent-face, respectively, against the scrambled-face preview. Differently from the previous comparison, it is now evident that seeing a face rather than a scramble-face stimulus in the periphery led to a completely different response pattern at the time of the new fixation, once the target face was foveated. While the fixation-related P1 did not differ between conditions, the following N1 component was dramatically influenced by the type of preview visible during the preceding fixation. In particular, we report a strong positivity at occipito-temporal channels, corresponding to a strong attenuation of the fixation-related N170 in the conditions in which a congruent or incongruent face was visible before the saccade. This effect was more pronounced over the right hemisphere, and a corresponding negative pole over central frontal regions, congruent with the activation pattern observed for the ERP response time-locked to stimulus onset (see section above).

This is especially clear by looking at the three waveforms for the electrodes PO7/PO8, whereby the incongruent- (green) and congruent- (pink) face preview showed a strong reduction in the post-saccadic evoked response at the time of the N170, i.e. a *preview positivity* effect, when compared to the scrambled-face preview (blue) (Figure 4B). Figure 4C visualizes the complete *p*-values matrix for the contrast between the congruent-face minus scrambled-face preview condition across the entire epoch. This plot confirms that the preview positivity began at around 160 ms and persisted up to about 300 ms after fixation onset. This was then followed by a later and weaker cluster of activation between around 360 to 420 ms, which shared a similar scalp topography as the initial N170 effect. For completeness, we also report the *p*-values matrix for the contrast between the incongruent-face minus scrambled-face preview condition (Figure 4D). The TFCE statistical matrix highlights how landing on a different face from the one that was available during the preview led to an almost identical pattern of activation over all the electrodes for the entire epoch as in the congruent preview condition.

## Discussion

Neural correlates of face and object recognition have traditionally been studied by flashing stimuli to the central visual field during prolonged visual fixation. In contrast, natural vision typically affords an extrafoveal preview of soon-to-be fixated items before they are brought into the fovea with a saccade. Here we show that the extrafoveal preview of a face stimulus leads to a dramatic reduction in the post-saccadic evoked response to that stimulus, compared to a control condition in which a meaningful preview was withheld by scrambling the spectral phase of the preview stimulus. This reduction occurred even though a face stimulus was always present after the saccade, with only the pre-saccadic preview differing between the conditions. Interestingly, this preview reduction was similar for trials in which the exact same face was present across the saccade (congruent-face condition) and when a different face was present after the saccade (incongruent-face condition). In particular, the N170 component, which is classically linked to the structural processing of faces, was substantially reduced in trials with a face preview than those with a scrambled preview. More generally, the fixation-related evoked response to the post-saccadic face was strongly attenuated, consistent with a “preview benefit” (i.e. reduction) in the evoked response as previously observed for visual words (Dimigen et al., 2012). These findings are consistent with claims that both visual and category-specific information about the saccadic target can influence post-saccadic processing of that stimulus (for review, see: Melcher and Colby, 2008; Melcher and Morrone, 2015).

A critical issue in the context of preview studies is whether the peripheral preview influences early or late ERP components (Edwards et al., 2017). Interestingly, in our data, a significant preview effect arose only after the end of the P1 component, around 160 ms after fixation onset. Current theories suggest that face processing involves several stages of neural processing which differ in terms of their feature-selectivity, neural substrate, and associated ERP components. In particular, the *occipital face area* has been implicated in processing parts of faces, such as eyes or mouth, and suggested to influence the P100 component (Pitcher et al., 2007; Sadeh et al., 2010), while the *fusiform face area* is associated with processing facial identity and thought to be reflected in N170 activity, typically associated with “structural encoding” (Halgren et al., 2000; Hoshiyama et al., 2003; Deffke et al., 2007; Gao and Wilson, 2013). From our results, the peripheral preview related modulation is therefore more consistent with processing of facial identity and structural encoding (Sadeh et al., 2010), rather than individual parts. The lack of any difference between congruent and incongruent previews is also consistent with processing at the level of the face representation rather than specific local features. This result is compatible with the task in which the participants were involved, that was a gender discrimination, suggesting that participants might have focused more on the global features of the stimuli presented rather than any specific face part. At the same time, the preview faces in our study were presented in the periphery as part of a bilateral pair of stimuli, and so future work is needed to investigate whether earlier components might be influenced when the competition between two stimuli is removed and only one face stimulus is presented in isolation in the periphery.

The morphology of the effects observed here also argues strongly against an interpretation of preview effects as merely the absence of surprise or a change in context, since such effects are more typically reflected in the later centroparietal P3 component (Sutton et al., 1965; Duncan-Johnson and Donchin, 1977; Donchin, 1981). Indeed we observed a persistence of positivity also in the later part of the fERP (as well as in the ERP during fixation), that in the context of face stimuli might be associated with processing of dynamic aspects of the face in the Superior Temporal Sulcus (Itier and Taylor, 2004; Sadeh et al., 2010; Dalrymple et al., 2011). Also, there was a face stimulus present after the saccade in every trial, meaning that the face target was not a surprise in the traditional sense.

It is important to note that the overall pattern of results is not consistent with hypotheses based on a primary role for spatial attention or for priming. Prior to saccade execution attention is allocated to the saccade target (Hoffman and Subramaniam, 1995; Deubel and Schneider, 1996; Zhao et al., 2012; Buonocore et al., 2017) and such allocation might be associated with an enhancement of the P1 and N1 components of the fixation-related ERP (see also: Eimer, 2000; Mohamed et al., 2009; Sreenivasan et al., 2009; Churches et al., 2010; Meyberg et al., 2015). Results from a number of paradigms have in fact shown that stimuli that might be relevant as peripheral previews can lead to an increased evoked response. For example, repeating a face leads to increased (i.e. more negative) ERP amplitudes in the time window following the peak of the N170 (N250 early repetition priming effect; Schweinberger et al., 1995) rather than the decreased amplitude seen here for trans-saccadic preview. On the other hand, prediction mechanisms would have led to a reduction in evoked responses. Indeed, our results are consistent with the prediction hypothesis and the idea that prediction and attention can be dissociated by looking at the direction of effects, with enhanced evoked activity for attention and attenuated responses for prediction (Kok et al., 2012; Spaak et al., 2016; de Lange et al., 2018). Nonetheless, we cannot completely rule out that the relatively stronger response in the scrambled-face condition might be at least partially modulated by the visual discrepancy between the foveated face stimulus compared to the scrambled stimulus seen in the extrafoveal visual field during the preview interval. According to this framework, the larger N1 for the scrambled-face preview condition would share some features with the visual mismatch negativity (Stefanics et al., 2014; Kornrumpf et al., 2016). One could then argue that in the scrambled-face preview condition (i.e. the condition where the prediction is not matched), the onset of the face stimulus might lead to an enhanced “mismatch” response compared to conditions where participants were seeing a face before and after the saccade (i.e. conditions where the prediction is matched). In this regard, the interpretation of the results would be compatible with the idea of an increase in amplitude for the “unpredicted” condition.

It is also important to note that the fixation-related ERPs elicited by faces brought to the fovea by the saccade showed a generally similar N170 component to that traditionally observed for a sudden face onsets in the fovea (Soto et al., 2018). Given that face perception in real life is typically in response to a face brought to the fovea from the periphery, rather than a face appearing out of nowhere, it was critically important to show the ecological validity of such category-specific components. In future work, it would be interesting to see whether the fixation-related N170 behaves in a similar way to the classic ERP component under various manipulations such as inversion (Bentin et al., 1996).

Finally, our results are in line with the pattern of results reported in the decoding study by Edwards and colleagues (Edwards et al., 2017) but also expand on their work in several ways. First, we could at least in part address the issue of whether the modulation of the fERP represents a preview benefit rather than a change-related cost, due to including a scrambled-face condition that was not part of their design. In their study, there was only a valid or invalid preview condition. Some reduction in decoding performance might have resulted, for example, from the added noise in the invalid preview condition due to a visual mismatch negativity-related signal. Another way in which their results might reflect an invalid cost (rather than validity benefit) is that the use of a decoding measure means that information about the invalid preview might still have been available well into the new fixation, with that information interfering with the decoding of the post-saccadic stimulus. As described above, there is evidence that visual processing during the first 50-100 ms of a new fixation is driven at least in part by pre-saccadic information, followed by a potentially shift to processing the new input at the current foveal position. Future work is needed to clarify the question of how pre-saccadic object-specific information influences the earliest stages of visual processing and how this might depend on task requirements (such as the gender task used here).

More generally, the current study provides a proof of concept for the usefulness of the fERP paradigm for studying trans-saccadic integration and visual stability. Follow-up studies can use this technique to investigate the mechanisms underlying trans-saccadic perception to distinguish between competing theories of how our impression of the visual world remains stable despite the retinal image changing so dramatically an average of three times per second.

